## Supplemental Fig S1-S8 Table S1-S6 for "Meioc-Piwil1 complexes regulate rRNA transcription for differentiation of spermatogonial stem cells"

**Supplemental Material**

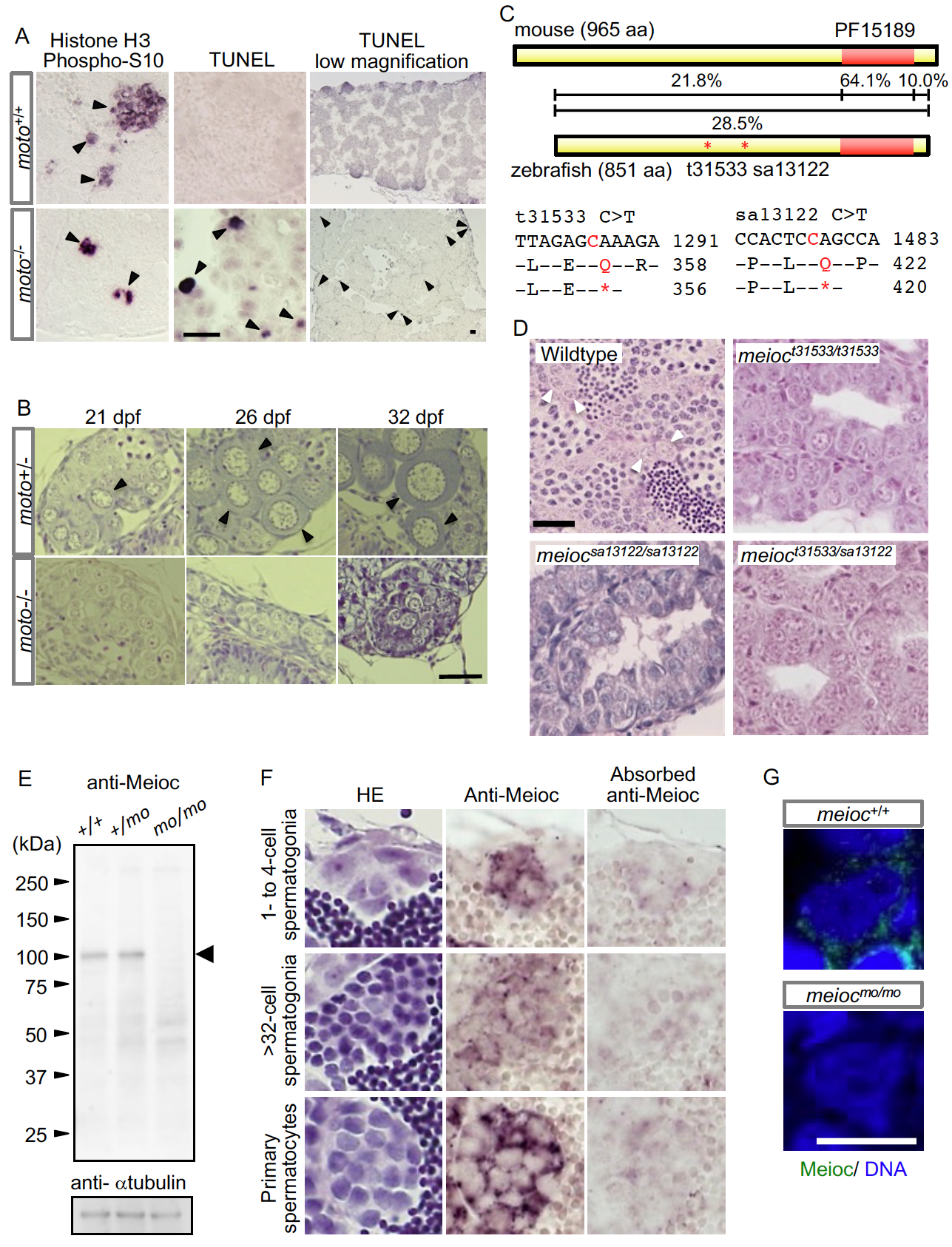

**Supplemental Figure S1. Phenotypes of testes and early gonads of the *moto^-/-^* mutant.**

(A**)** PH3 and TUNEL assays of wild-type and *moto^-/-^* testes. PH3-positive spermatogonia were detected in *moto^+/+^* and *moto^-/-^* testes. TUNEL-positive cells were barely detected in *moto^+/+^* testes, while a considerable number of cells were detected in *moto^-/-^* testes. Black arrowheads indicate positive spermatogonia.

**(**B**)** Histology of early gonads of *moto^+/-^* and *moto^-/-^*. Growing oocytes (arrowheads) were observed in heterozygous larvae but not in mutant larvae. Developmental stages are indicated by days post-fertilization (dpf). Scale bar, 20 µm.

(C) Comparison of Meioc mouse and zebrafish protein sequences. The sequence of zebrafish Meioc (ENSDARG00000090664) was aligned to that of mouse Meioc (ENSMUSG00000051455). Alignment analyses were performed using EMBOSS NEEDLE, an online software program (<https://www.ebi.ac.uk/Tools/psa/emboss_needle/>). A coiled-coil domain composed of four helixes, PF15189 (previously DUF4582), is conserved in animals. The two mutations predicted to disrupt the zebrafish *meioc* gene are shown, and the approximate position of the premature stop codon is indicated relative to the total length of the Meioc protein.

(D) HE-stained sections of wild-type testes with all stages of spermatogonia (arrowheads), spermatocytes and spermatozoa present; the *meioc^t31533^* mutant, an additional nonsense allele *meioc^sa13122^,* and transheterzygote *meioc^t31533/sa13122^* exhibited testes containing single spermatogonia and spermatogonia in small clusters. Scale bar: 20 µm.

(E) Western blot analysis of *meioc^+/+^, meioc^+/mo^* and *meioc^mo/mo^* testis extracts.

(F) Immunostaining of zebrafish wild-type testes. Absorbed anti-Meioc IgG was treated with Meioc recombinant protein. Scale bar: 10 µm.

(G) Immunostaining of *meioc^mo/mo^* single spermatogonia. Note that Meioc was not detected in *meioc^mo/mo^* spermatogonia. Scale bar, 10 µm.

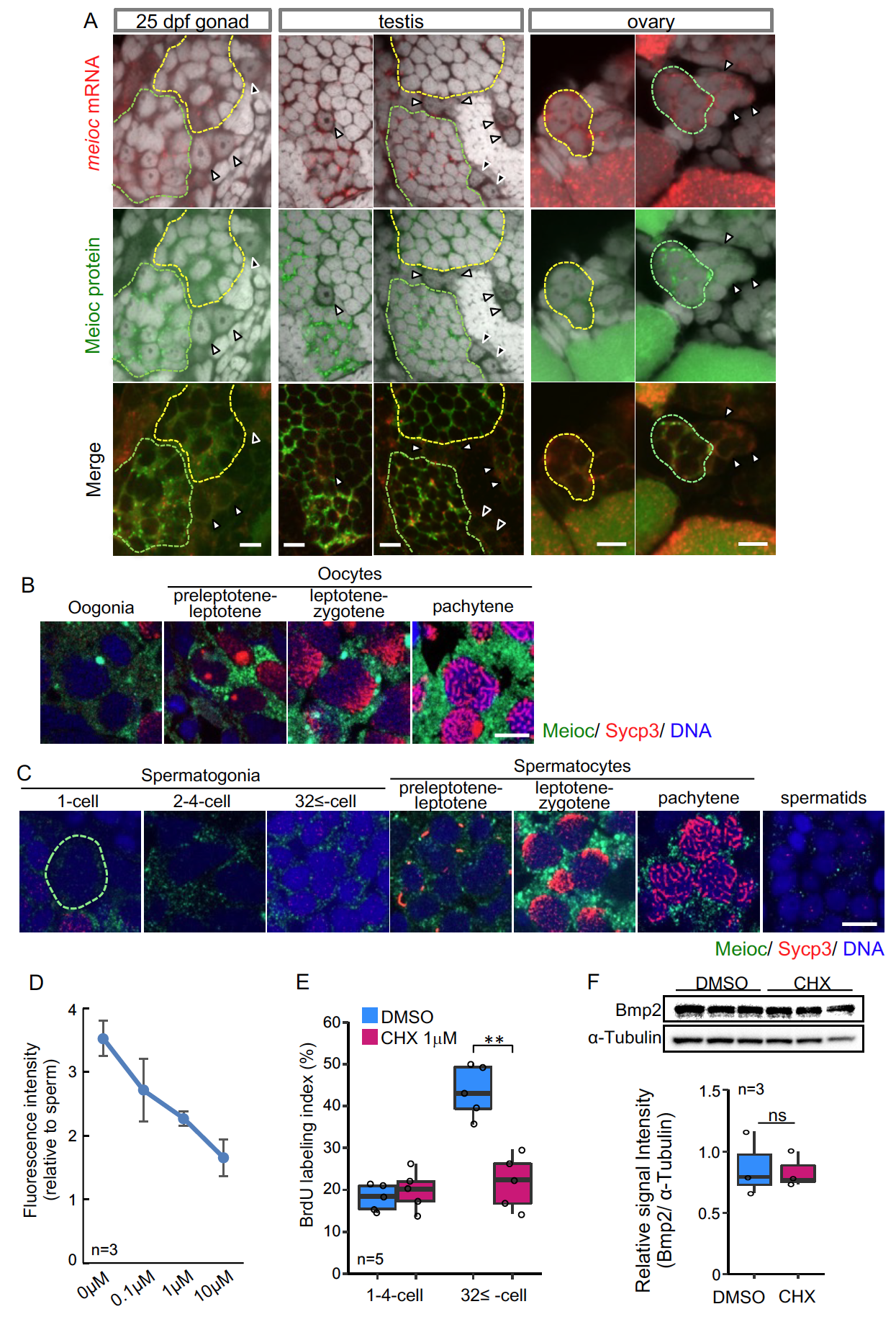

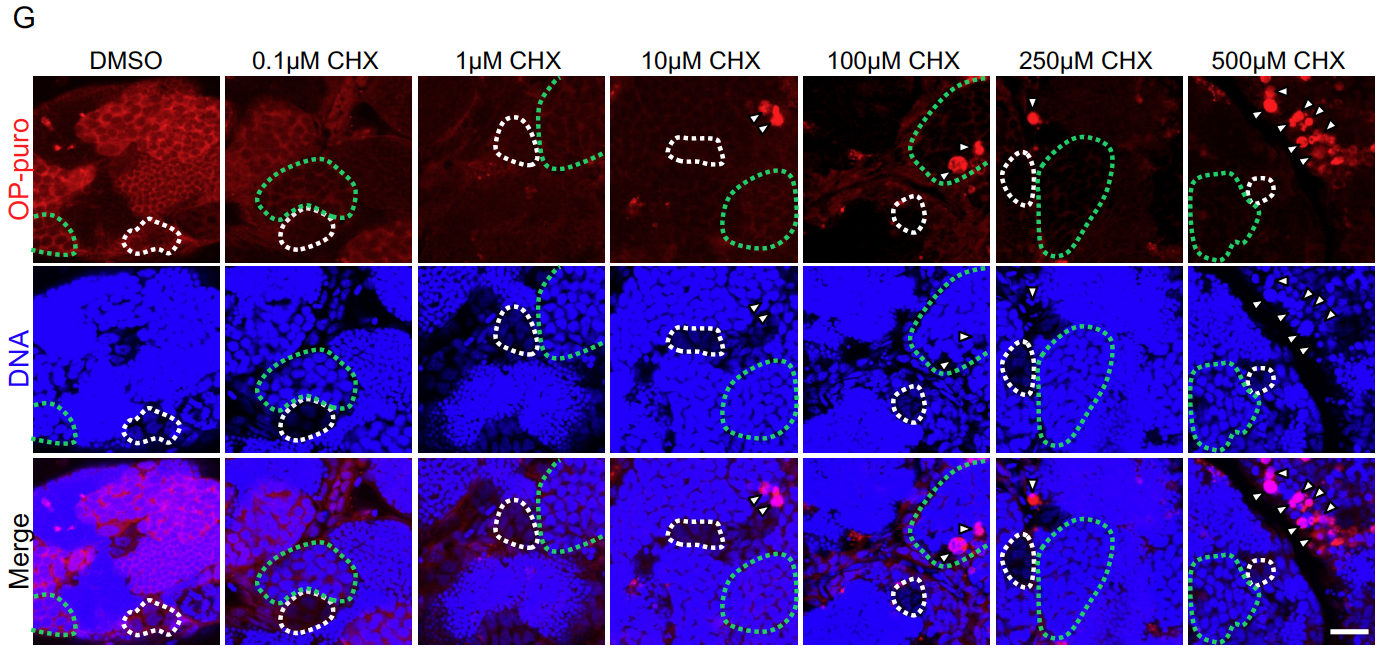

**Supplemental Figure S2. Expression patterns of Meioc in wild-type germ cells and effect of cyclohexmide on spermatogonia.**

(A) Expression patterns of *meioc* mRNA and Meioc protein in gonads at 25 dpf and adult testes and ovaries. Black arrowheads indicate cells that express neither mRNA nor protein, and white arrowheads indicate cells that have the mRNA signals and weak protein signals. Yellow dotted lines indicate cysts containing cells positive for both mRNA and diffuse protein signals, including small or weak granules, and green dotted lines indicate cysts containing cells with mRNA signals and bright protein signals, including large granules. Scale bar, 10 µm.

(B) Double staining of ovary at 25 dpf with anti-Meioc antibody (green) and anti-Sycp3 antibody (red). Staging of oocytes was defined by the patterns of Sycp3. Scale bar, 5µm.

(C) Double staining of wild-type testis with anti-Meioc antibody (green) and anti-Sycp3 antibody (red). Staging of spermatogonia was defined by the number of cells in the cyst, and staging of spermatocytes was defined by the patterns of Sycp3. Scale bar, 5 µm.

(D) Dose-dependent protein synthesis by cycloheximide (CHX) in late spermatogonia of the cultured testis.

(E) Effect of CHX (1 µM) on BrdU incorporation of cysts of 1-4-cell and 32≤-cell spermatogonia in testis organ culture.

(F) Western blot analysis of Bmp2 and α-Tubulin (upper panels) and quantification of Bmp2 (lower panel) in the cultured feeder cells with and without 0.2 μM cycloheximide.

(G) Toxicity of CHX in cultured testis fragments. CHX was treated with testicular fragments for 48 hours at various concentrations. Note that abnormal strong OP-puro signals indicated by arrowheads including nuclei were detected in cells at 10 μM or more. White dotted line: cysts containing 1-4-cell spermatogonia. Green dotted line: cysts containing spermatocytes. Scale bar, 20 µm.

For each graph, data were analyzed by Student’s t test: **p < 0.01. ns: no significant difference,

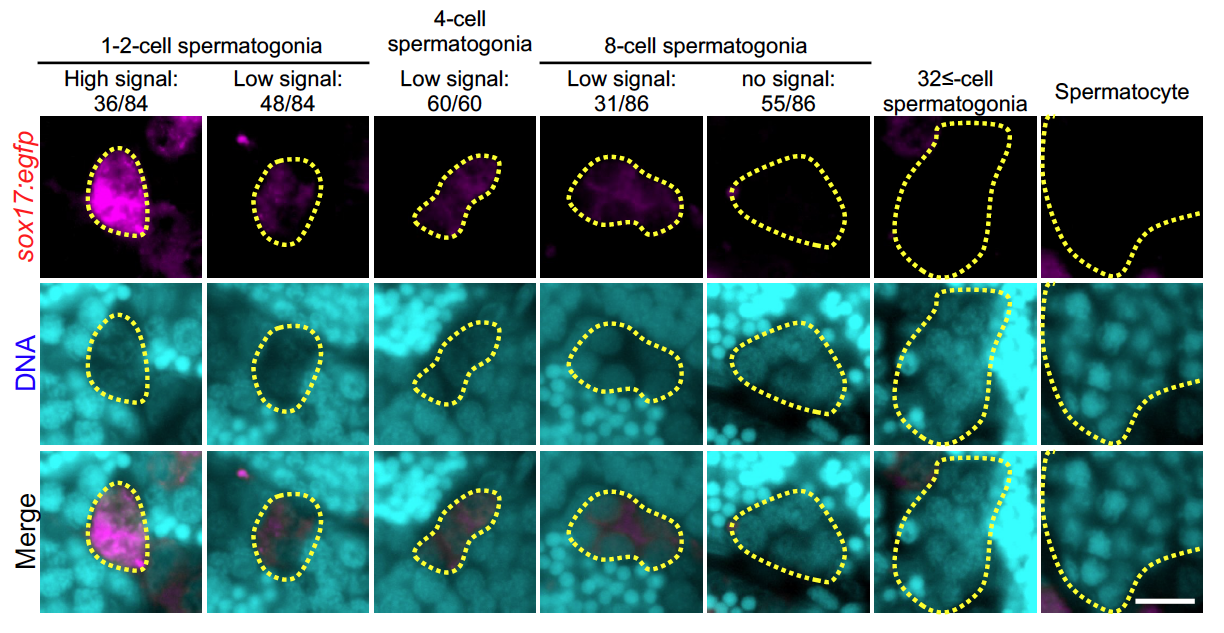

**Supplemental Figure S3. Expression pattern of EGFP in the *sox17:egfp* transgenic testis.**

Staining of *sox17:egfp* testis sections with anti-GFP antibody (magenta) and DAPI (cyan). Note that EGFP was strongly expressed in the 1-2-cell stage spermatogonia and weakly expressed in the 4-8-cell spermatogonia, and started to fade at the 8-cell spermatogonia. Stem cell function of *sox17:egfp* spermatogonia was already confirmed by transplantation experiment (Kawasaki et al., 2016). Dotted lines indicate cysts containing the cell types indicated above in the image.

Scale bar, 10 µm.

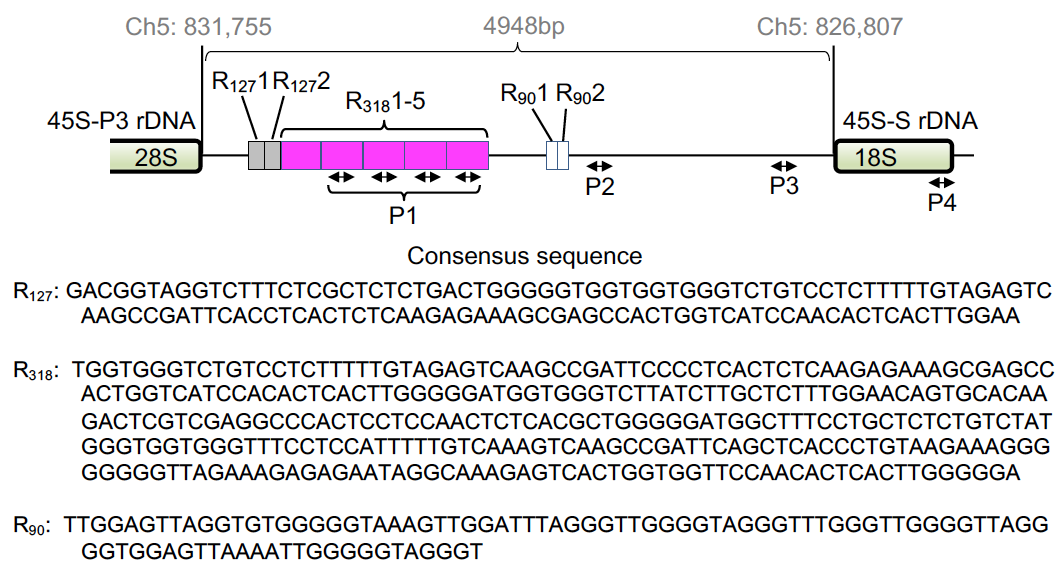

**Supplemental Figure S4. Schema of the IGS (chromosome 5: 826807-831755 reverse strand) and the 120, 318, and 90-bp tandem repeat sequences.** Two-way Arrows indicate position of primers used in Chip-qPCR analysis (Figure 6 F and G). See also Table S4.

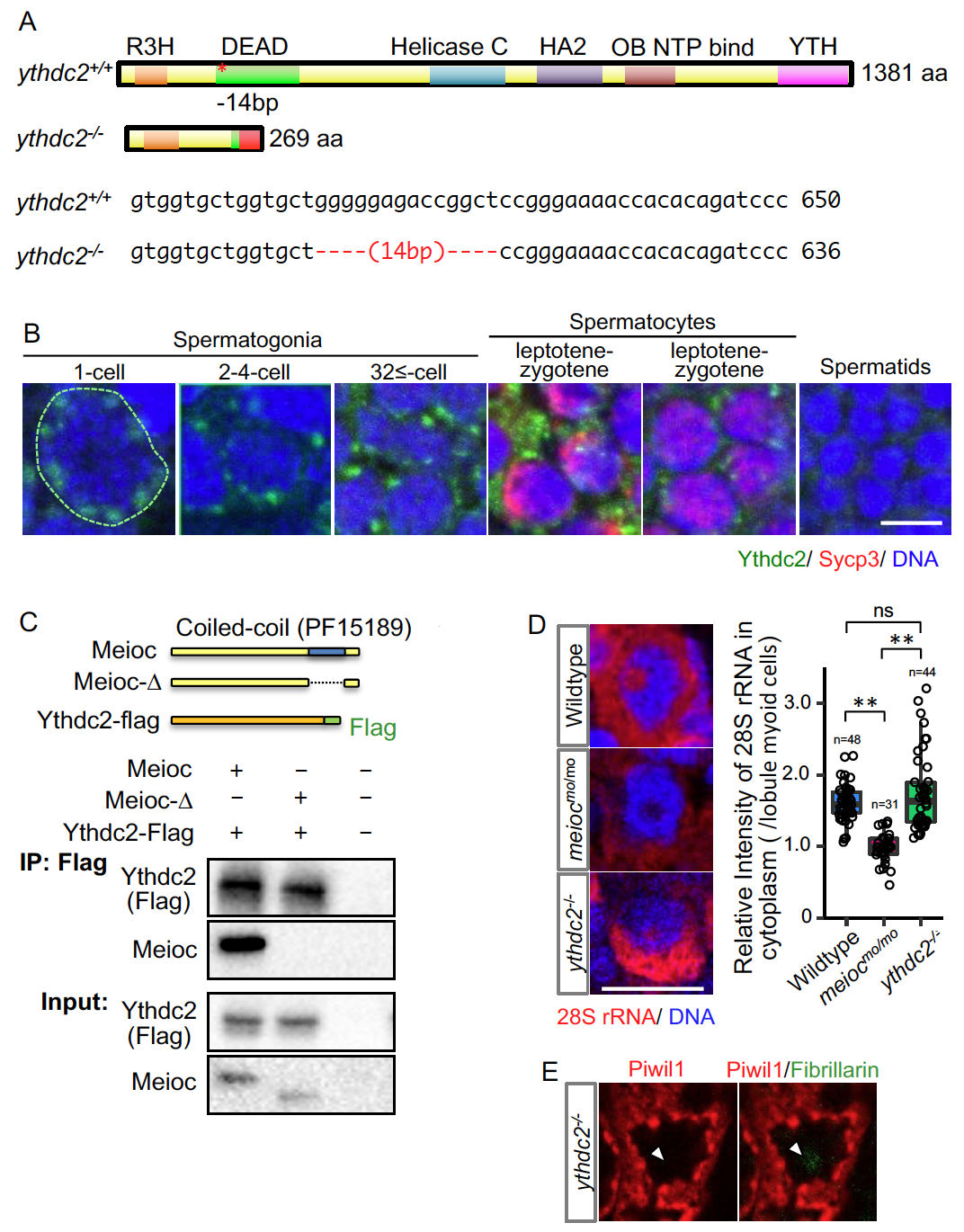

**Supplemental Figure S5.** **Mutations of the *ythdc2^-/-^*.**

(A) Zebrafish Ythdc2 protein structure and mutation sequence (*) in the *ythdc2* KO zebrafish.

(B) Double staining of testis with anti-Ythdc2 antibody (green) and anti-Sycp3 antibody (red). Staging of spermatogonia was defined by the number of cells in the cyst, and staging of spermatocytes was defined by the patterns of Sycp3. Scale bar, 5µm.

(C) Pull down assay using full-length or coiled-coil domain deleted Meioc and Flag-tagged Ythdc2. Schema: conditions of coexpression of each protein in HEK-293 cells used for the pull-down assay.

(D) In situ hybridization of 28S rRNA (left panels) and quantification of the 28S rRNA signal intensities (right panels) in wild-type, *meioc^mo/mo^* and *ythdc2^-/-^* 1-2-cell spermatogonia. The cytoplasmic signal intensities were normalized to the myoid cell cytoplasm.

(E) Immunostaining against Piwil1 (red) and fibrillarin (green) in *ythdc2^-/-^*spermatogonia. Arrowheads; undetectable Piwil1 signals in Fibrillarin (green) positive nucleoli under the normal sensitivity imaging of Piwil1.

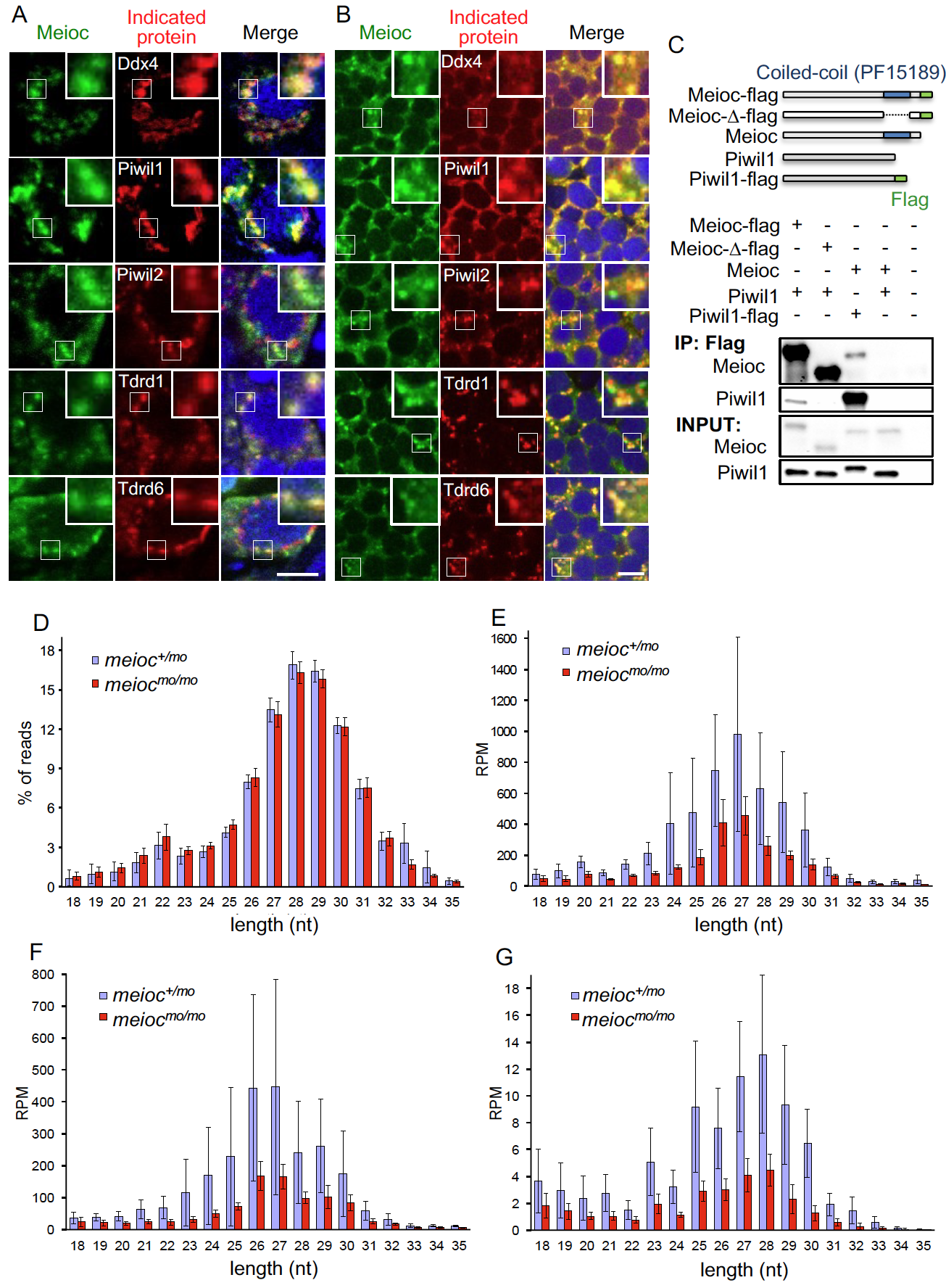

**Supplemental Figure S6. Immunostaining of Meioc with germ granule components and piRNA profiles in the *meioc^mo/mo^* testis.**

(A, B) Immunostaining of Meioc (green) and Ddx4, Piwil1, Piwil2, Tdrd1 and Tdrd6 (red) in wild-type 1-2-cell spermatogonia (A) and spermatocytes (B). Insets show magnified signals of each spot indicated by a white square. Blue in right panels = DAPI. Scale bar; 5 μm.

(C) Pull down assay using Flag-tagged full-length Meioc, coiled-coil domain (PF15189)-deleted Meioc and Piwil1. Schema: conditions of coexpression of each protein in HEK-293 cells used for the pull-down assay.

(D-G) Small RNAs from the testes of individual animals with either the *meioc^+/mo^* and *meioc^mo/mo^* genotypes were sequenced. Data from the individuals were then pooled for each genotype to assess the global effect of Meioc on small RNA production (mean +/- s.d. of 6 *meioc^+/mo^* and 5 *meioc^mo/mo^* testes). Length profiles of the relative abundance of all 18-35 nt mapped reads (D), as well as only the 28S rRNA (E), 18S rRNA (F) and R2 transposon (G) mapped reads. Significant increases in the small RNAs derived from 18S, 28S rRNA and R2 transposon were not detected in *meioc* mutant testes, although Piwil1 was ectopically localized in nucleoli in the mutant spermatogonia. RPM: Reads Per Million.

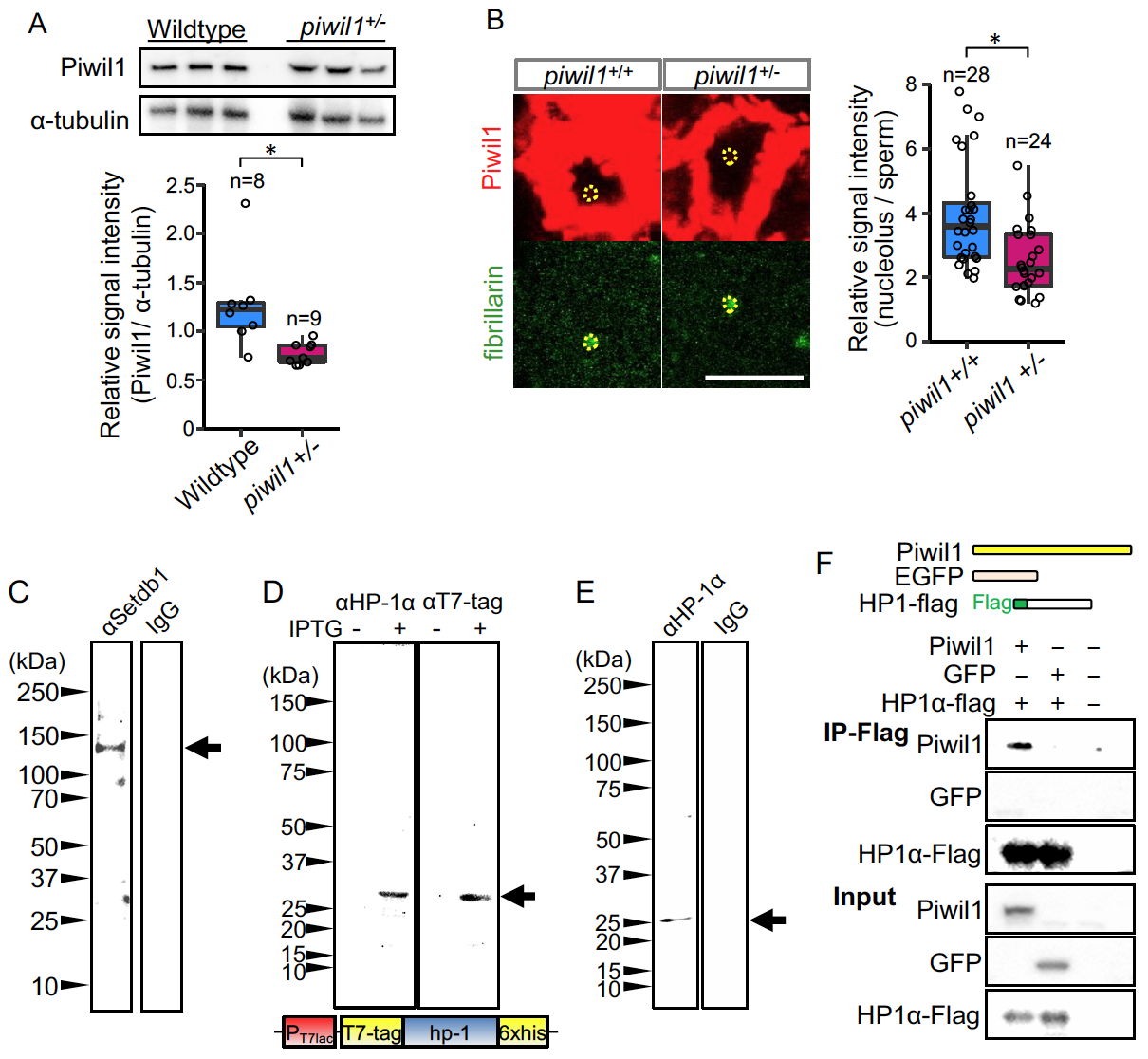

**Supplemental Figure S7. Reduction of Piwil1 in *piwil1^+/-^* and interaction of Piwil1 with Setdb1 and HP1α but not Dnmt3a.**

(A) Western blot analysis of Piwil1 and α-Tubulin (upper panels) and quantification of Piwil1 (lower panel) in *piwil1^+/+^* and *piwil1^+/-^* testes.

(B) Immunostaining against Piwil1 and fibrillarin (left panels) and quantification of nucleolar Piwil1 signal intensities (right panel) in *piwil1^+/+^* and *piwil1^+/-^* spermatogonia (1-2-cell cysts). Yellow dotted lines: nucleoli.

(C) Western blot analysis of wild-type testis extracts using anti-mouse Setdb1 antibody. Arrow: Setdb1.

(D) Western blot analysis of zebrafish HP1α expressed from bacterial expression vectors using anti-human HP1α antibody and anti-T7-tag antibody. IPTG was added for culture of host cells (BL21) to induce expression of HP1α with T7-tag. Schema: construction of expression vector (pet21a). Arrow: HP1α fusion protein.

(E) Western blot analysis of wild-type testis extracts using anti-human HP1α antibody. Arrow: HP1α.

(F) Pull down assay using Flag-tagged HP1α and Piwil1 or EGFP. Schema: conditions of coexpression of each protein in HEK-293 cells used for the pull-down assay.
For each graph, data were analyzed by Student’s t test: *p < 0.05.

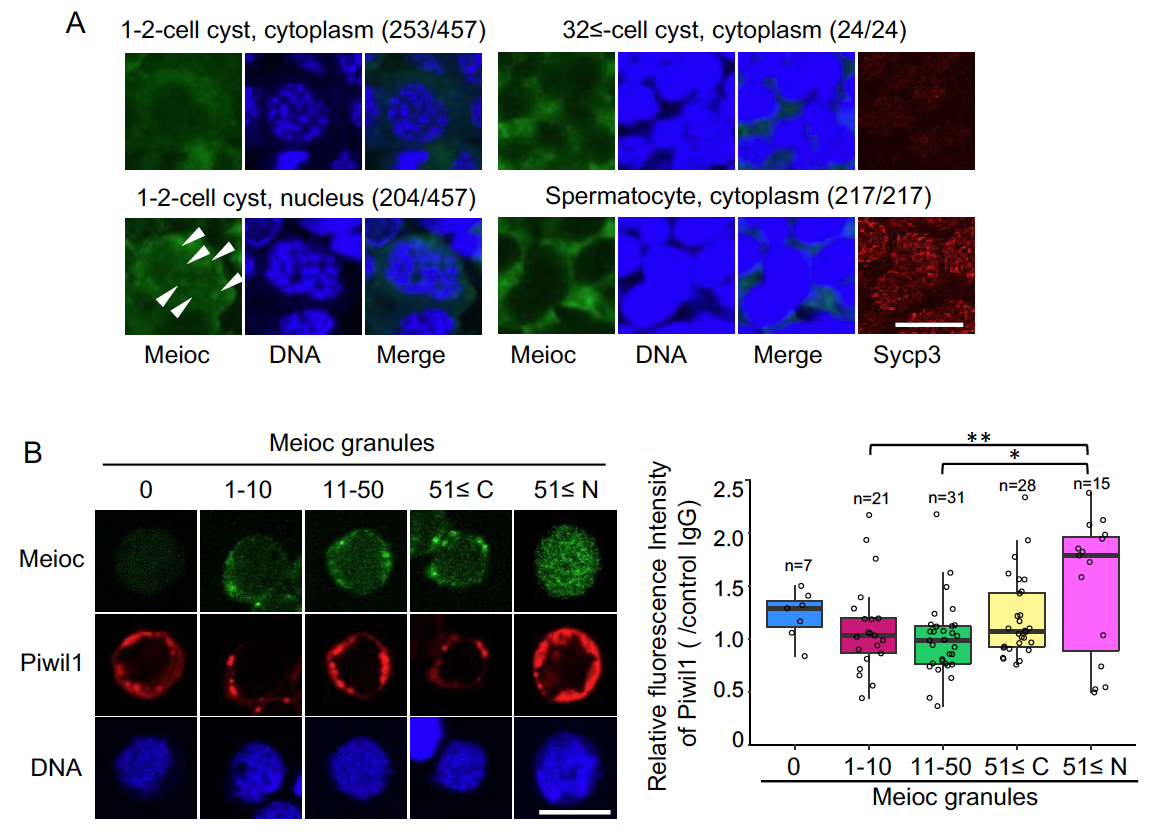

**Supplemental Figure S8. Intracellular localization of Meioc in spermatogenic cells and Piwil1 expression levels in isolated isolated *sox17::egfp* spermatogonia.**

(A) Meioc localization in 1-2-cell and 32≤-cell cyst spermatogonia and spermatocytes in testis sections. To distinguish spermatocytes from 32≤-cell spermatogonia, Sycp3 was stained. The number in parentheses is the corresponding cells in total.

(B) Expression pattern of Piwil1 in isolated *sox17::egfp* spermatogonia, based on the amount of Meioc granules and the localization. Right panels are intensities of Piwil1 for each class of the purified wild-type *sox17::egfp* spermatogonia. 51≤ C and N; 51≤ cytoplasmic and nuclear Meioc granules, respectively. *p < 0.05, **p < 0.01, n: number of analyzed spermatogonia. Scale bars: 10 µm.

**Supplemental Table S1.** **Sex ratio of *meioc* mutants and heterozygotes.**

|  | Female | Male |
| --- | --- | --- |
| *meioc^mo/mo^* | 0 | 24 |
| *meioc^+/mo^* | 23 | 14 |
| *meioc^+/+^* | 13 | 13 |

**Supplemental Table S2.** **LC/MS/MS for the immunoprecipitate (IP) of Meioc with lysate of a wild-type testis.**

Proteins in anti-Meioc IP sample with at least 3-fold enrichment in normal testes compared with control IgG IP sample, as detected by mass spectrometry. Uniprot ID are shown in Accession. Proteins were classified with GO-terms (http://amigo.geneontology.org). The molecular weight, number of peptides identified by spectrometry, and coverage are provided. Meioc protein is highlighted as red, and known mouse MEIOC partner YTHDC2 is indicated by bold.

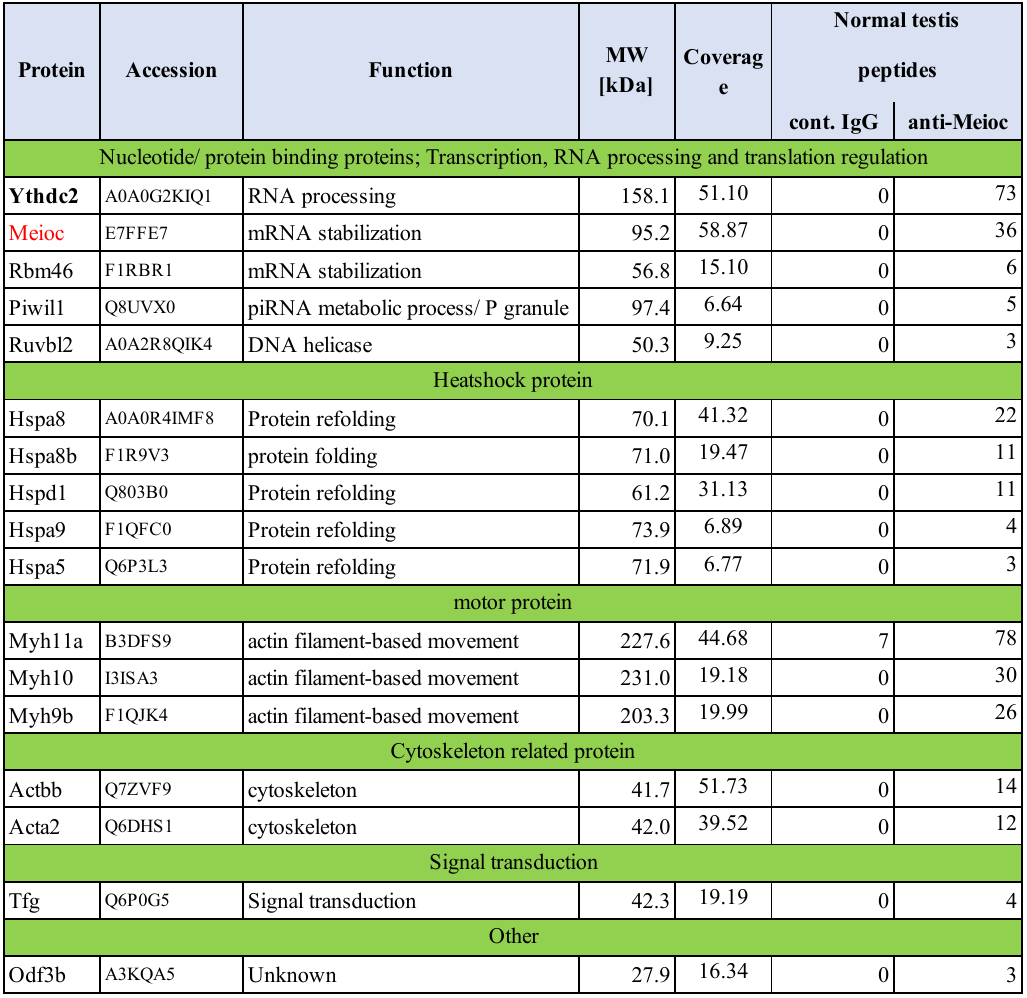

**Supplemental Table S3.** **LC/MS/MS for the immunoprecipitate (IP) of Meioc with lysate of a hyperplasia testis in which GSCs accumulate.**

Proteins in anti-Meioc IP sample with at least 3-fold enrichment in normal testes compared with control IgG IP sample, as detected by mass spectrometry. NCBI protein ID or Uniprot ID are shown in Accession. Proteins were classified with GO-terms (http://amigo.geneontology.org). The molecular weight and number of peptides identified by spectrometry, and coverage are provided. Meioc protein is highlighted as red. *:protein recorded in RNA Granule Database as processing body (P-body) protein, **:protein recorded in RNA Granule Database as stress granule protein (Youn et al. 2018).

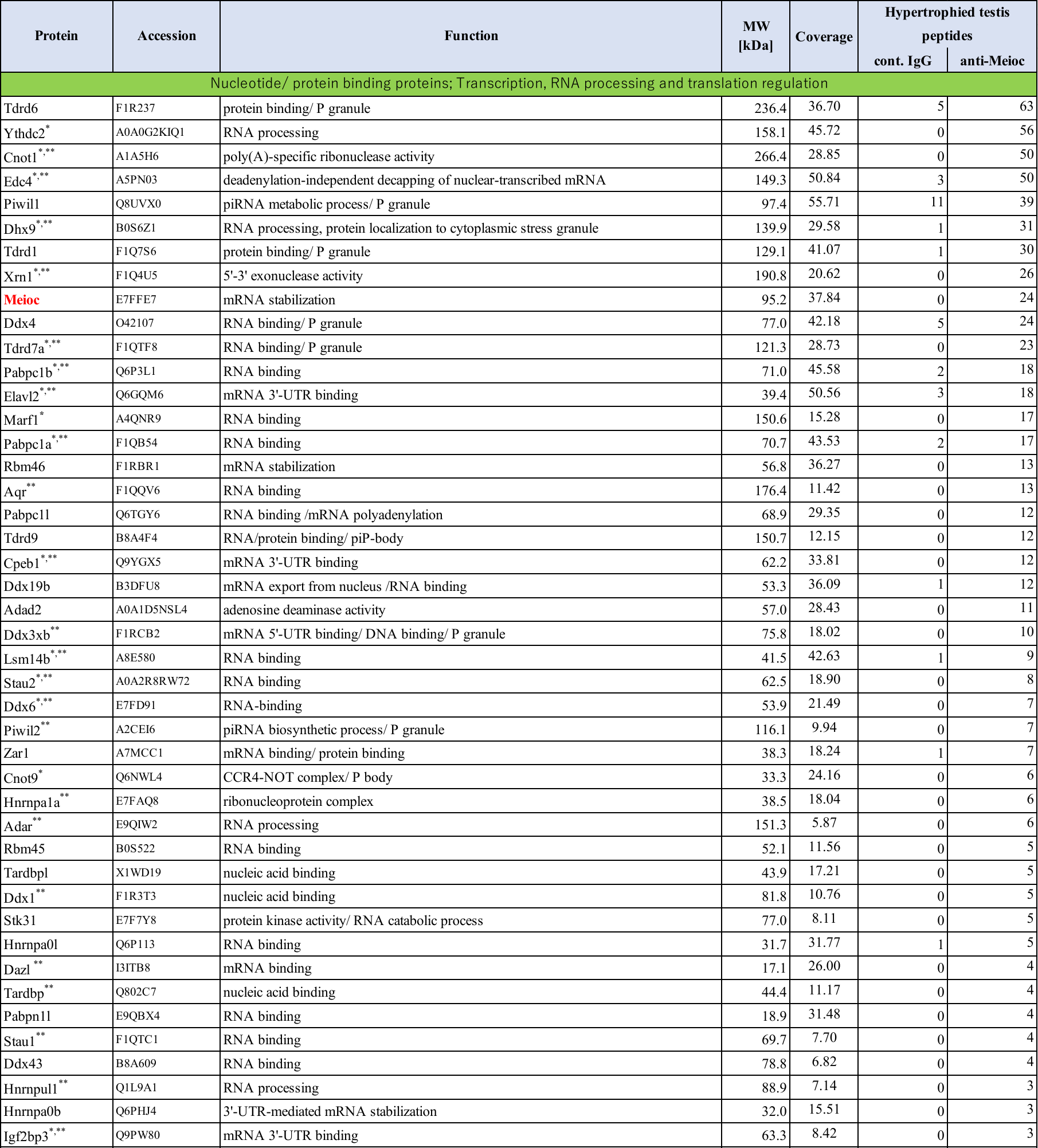

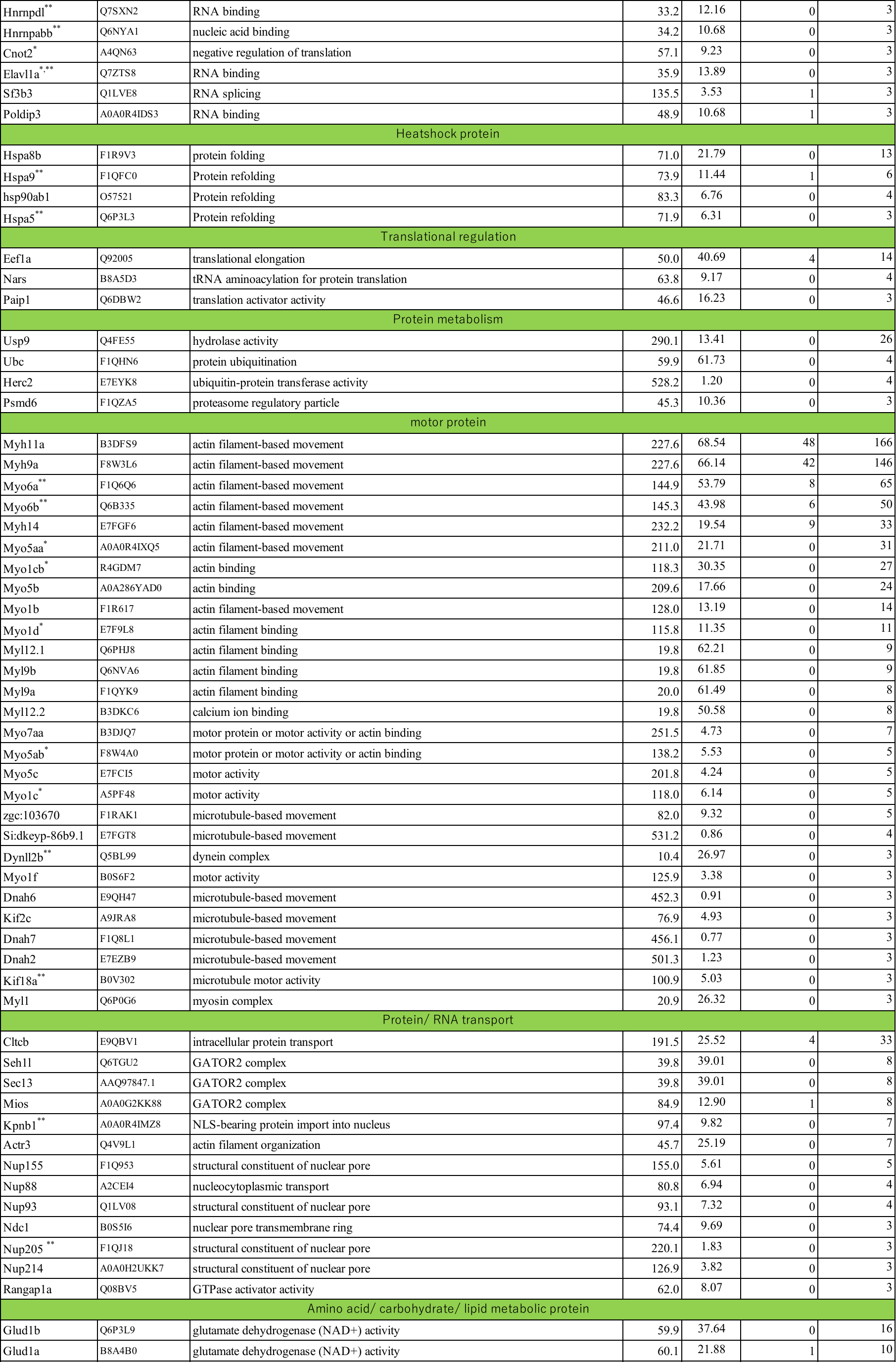

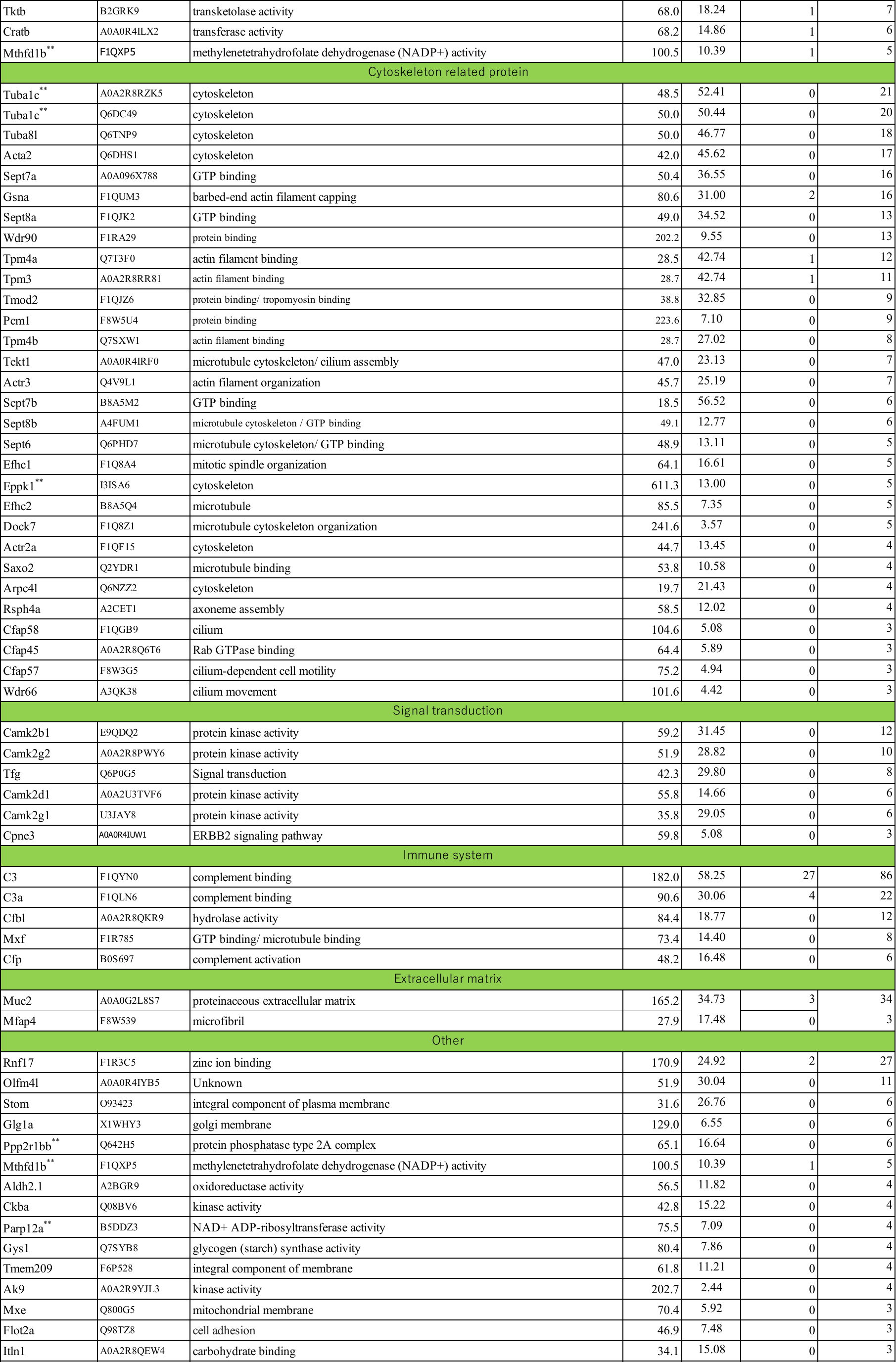

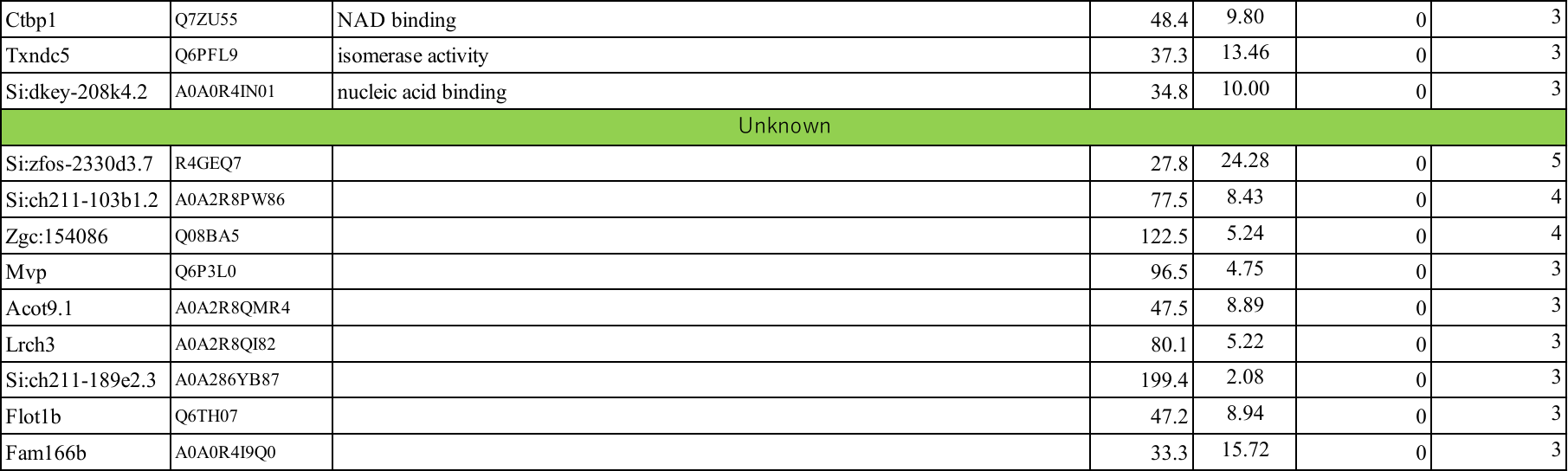

**Supplemental Table S4.** **Loci of tandem repeats in IGS region of 45S-S rDNA.**

| Name | Location | Start* | End* | Length |
| --- | --- | --- | --- | --- |
| R_127_1 | Chromosome 5 | 831,438 | 831,312 | 127 |
| R_127_2 | Chromosome 5 | 831,311 | 831,194 | 118 |
| R_318_1 | Chromosome 5 | 831,275 | 830,963 | 313 |
| R_318_2 | Chromosome 5 | 830,962 | 830,638 | 325 |
| R_318_3 | Chromosome 5 | 830,637 | 830,316 | 322 |
| R_318_4 | Chromosome 5 | 830,315 | 829,988 | 328 |
| R_318_5 | Chromosome 5 | 829,987 | 829,661 | 327 |
| R_90_1 | Chromosome 5 | 829,067 | 828,977 | 91 |
| R_90_2 | Chromosome 5 | 828,977 | 828,886 | 92 |

*: position in the latest release of reference genome assembly, GRCz11.

**Supplemental Table S5. Oligonucleotide primers and sgRNA used in this study**

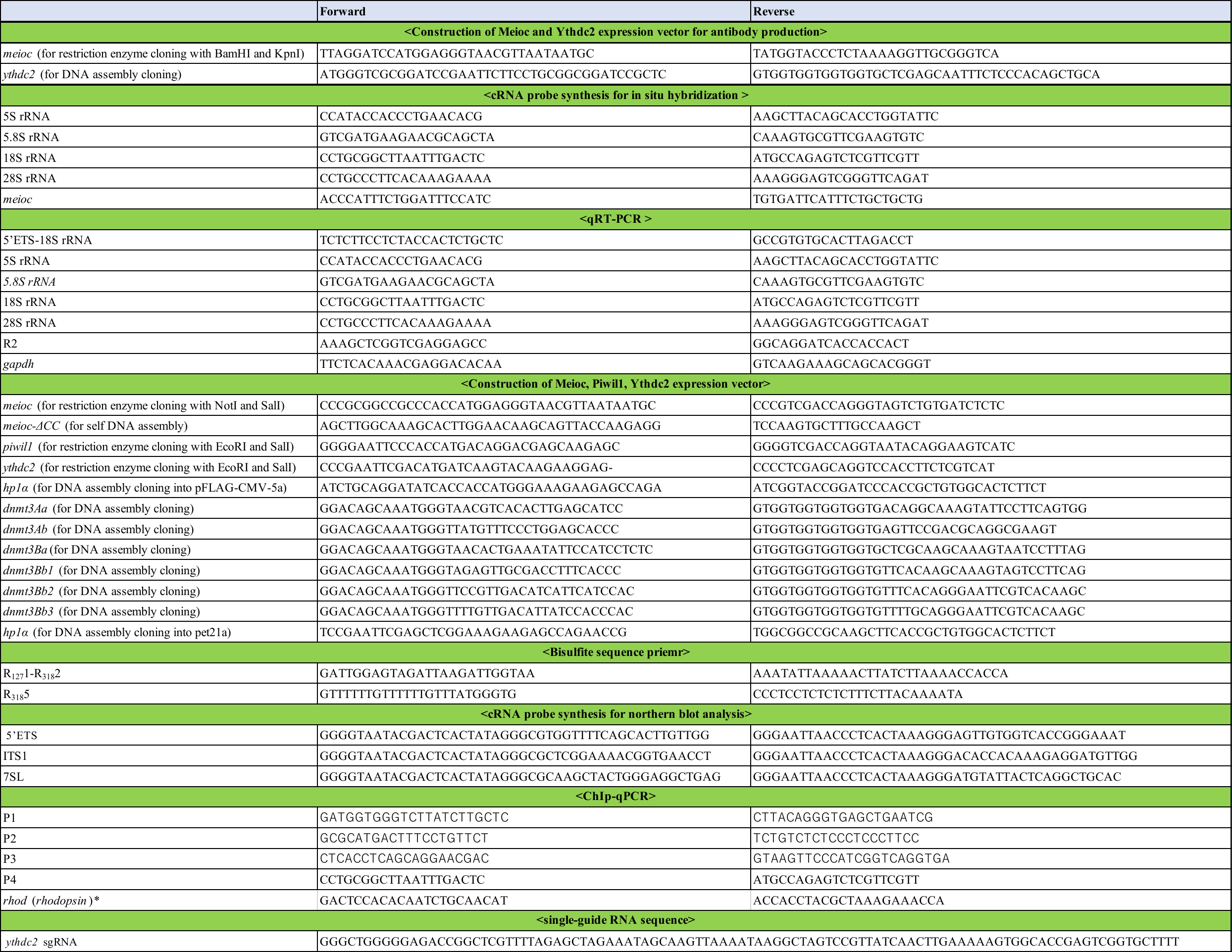

*:Primer set to amplify upstream of *rhodopsin* locus (Morley et al. 2009).

**Supplemental Table S6.** **Antibodies and reagents used in this study**

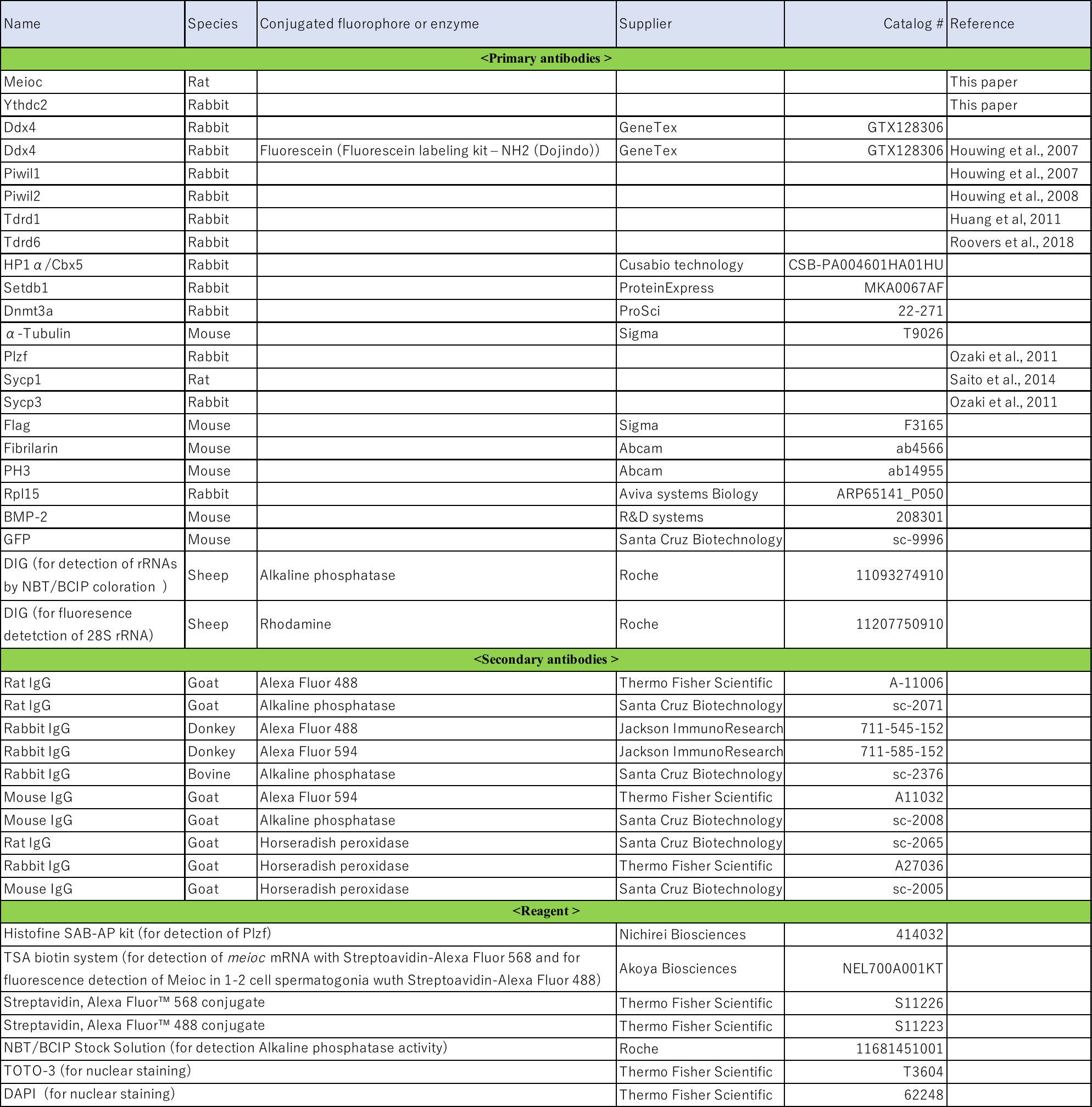
